# Traumatic brain injury alters hepatic gluconeogenic metabolism assessed using hyperpolarized pyruvate

**DOI:** 10.64898/2026.08.29.747003

**Authors:** Zohreh Erfani, Baljeet Seniwal, Erik J. Plautz, Joseph Park, Sasanka Wathukara Dewage, Sung-Han Lin, Shawn C. Burgess, Eunsook S. Jin, Jae Mo Park

**Author notes:** Correspondence to: Jae Mo Park, Ph.D. 5323 Harry Hines Blvd. Dallas, Texas, United States, 75390-8568, /. Zohreh Erfani and Baljeet Seniwal equally contributed to the work. **Conflicts of Interest**: Nothing to report.

## Abstract

**Background:** Acute phase response is an early immunometabolic response to brain injuries, primarily coordinated by the liver via the activation of acute phase proteins. These immune responses can be both beneficial, promoting tissue repair, and detrimental, exacerbating neurological deficits, if not properly controlled. Despite the central role of the liver in immunometabolism, how hepatic metabolism dynamically adapts to traumatic brain injury remains under explored, primarily due to limited liver-specific modalities that can assess metabolic pathways *in vivo*. ^13^C MRI utilizing hyperpolarized ^13^C-pyruvate can assess key regulatory enzyme activities in hepatic metabolism.

**Methods:** Rats with controlled cortical impact were studied *in vivo* using hyperpolarized [1-^13^C]pyruvate and [2-^13^C]pyruvate under fed and fasted conditions 3-4 days after injury. Hyperpolarized ^13^C products, including [^13^C]bicarbonate from [1-^13^C]pyruvate and [5-^13^C]glutamate, [1-^13^C]acetyl-L-carnitine, and [2-^13^C]phosphoenolpyruvate from [2-^13^C]pyruvate, were evaluated to assess mitochondrial and gluconeogenic metabolism. In parallel, liver tissues were collected following [U-^13^C_3_]pyruvate injection for NMR isotopomer analysis of phosphoenolpyruvate, glucose, and glutamate.

**Results:** While no metabolic differences were detected under fed condition, [^13^C]bicarbonate and [2-^13^C]phosphoenolpyruvate increased after brain injury under fasted condition, indicating an upregulation of the hepatic gluconeogenic pathway after injury. ^13^C NMR of liver tissue extracts from injured rats showed an elevated [2,3-^13^C_2_]glutamate-to-[4,5-^13^C_2_]glutamate ratio and increased ^13^C-labeling in phosphoenolpyruvate than controls, confirming enhanced hepatic gluconeogenic pathway.

**Conclusion:** This study demonstrates that hepatic acute phase response to brain injuries can be monitored *in vivo* by hyperpolarized pyruvate, which may be further utilized for longitudinal immunometabolic evaluation of the liver during pathogenesis and therapeutic interventions.

## INTRODUCTION

Traumatic brain injury (TBI) is a major global health issue, affecting all age groups and contributing to mortality and long-term disability.^1,2^ The pathology of TBI is largely characterized by two distinct phases: an initial mechanical insult followed by a secondary injury phase driven by neuroinflammation and metabolic disruption.^3^ Beyond direct neurological damage, TBI triggers systemic metabolic and immune alterations throughout the body.^4^ These extracranial effects include altered sympathetic and hormonal stress responses, metabolic disturbances, and multiorgan dysfunction involving the heart, liver, and kidneys.^5^

The immune system is a key mediator of the interconnection between the brain and peripheral organs, dynamically balancing pro- and anti-inflammatory processes that can either aid recovery or exacerbate damage.^6^ These immune responses can influence long-term neurological outcomes, such as neurodegeneration and chronic inflammation. Situated at the intersection of immune function and systemic metabolism, the liver serves as the largest reservoir of resident macrophages and coordinates the systemic response to brain injury through the production of cytokines and chemokines that regulate neuroinflammation.^7–9^

During the secondary injury, trauma signals activate cytokine production via vagal afferents and blood-borne mediators.^7,10^ Inflammatory mediators released from damaged brain tissue, particularly cytokines such as TNF-α, IL-1β, and IL-6, enter the circulation and activate hepatic signaling pathways. This induces a rapid acute phase response (APR), characterized by NF-κB-mediated transcriptional activation of acute phase proteins, including C-reactive protein, along with the release of additional cytokines and chemokines into the systemic circulation.^7,11^ Notably, among these mediators, CCL2 (MCP-1) is rapidly elevated after brain injury and promotes the recruitment of immune cells to both the brain and peripheral organs.^12^

In parallel, hepatic metabolism adapts to meet the increased nutritional demands of the injured brain, providing essential substrates like glucose and lactate for tissue repair and metabolic homeostasis. As a result, hyperglycemia is commonly observed in TBI patients. Managing this hepatic immunometabolic adaptation is critical since post-injury hyperglycemia is associated with poor clinical outcomes and mortality.^13–15^ Indeed, several studies reported neuroprotective effects of therapeutic strategies targeting hepatic metabolism and glucose homeostasis during TBI recovery, including insulin therapy, anti-diabetic drugs, soluble epoxide hydrolase inhibitors, and the ketogenic diet.^16–19^

Despite the critical role of the liver in modulating systemic post-TBI outcomes, how hepatic metabolism adapts to TBI *in vivo* remains poorly understood, and noninvasive tools capable of monitoring liver-specific responses are currently lacking. Hyperpolarized (HP) ^13^C magnetic resonance spectroscopy (MRS) overcomes this limitation by transiently enhancing ^13^C signals more than 10,000-fold and capturing real-time *in vivo* metabolism. HP [1-^13^C]pyruvate can be directly metabolized to [1-^13^C]lactate, [1-^13^C]alanine, and [^13^C]bicarbonate via lactate dehydrogenase (LDH), alanine aminotransferase (ALT), and pyruvate dehydrogenase (PDH), respectively.^20^ Using HP [1-^13^C]pyruvate, *in vivo* imaging of impaired cerebral PDH flux has been demonstrated during the acute phase of TBI by detecting [^13^C]bicarbonate in both animals^21,22^ and patients.^23^ In the liver, however, [^13^C]bicarbonate can be generated predominantly through phosphoenolpyruvate carboxykinase (PEPCK)-mediated decarboxylation following pyruvate carboxylase (PC) flux and isotopic scrambling between oxaloacetate and fumarate, particularly under fasting when PDH is suppressed.^24–26^ HP [2-^13^C]pyruvate provides more direct insight without this ambiguity between oxidative and anaplerotic/gluconeogenic pathways by detecting [5-^13^C]glutamate from PDH-TCA cycle flux and [2-^13^C]phosphoenolpyruvate (PEP) from PC-PEPCK flux, respectively.

In this study, we investigate the alteration of hepatic metabolism in response to TBI using *in vivo* HP [1-^13^C]pyruvate and HP [2-^13^C]pyruvate, complemented by *ex vivo* NMR isotopomer analysis of tissue extracts.

## METHODS

### Animal Preparation and TBI Model

All animal procedures followed the Guide for Care and Use of Laboratory Animals of US National Research Council and were approved by the University of Texas Southwestern Medical Center Institutional Animal Care and Use Committee (Protocol #:2017-101802) in accordance with the Animal Research: Reporting In Vivo Experiments (ARRIVE) guidelines.

Male Sprague Dawley rats (200-250 g, n = 36) were subjected to controlled cortical impact (CCI) injury,^27^ sham surgery, or used as healthy controls.^21^ Rats were anesthetized with isoflurane (induction: 3.5-4.5%, maintenance: 2.0-2.5%, in a 70% N_2_O, 30 % O_2_) and secured in a stereotaxic device (David Kopf Instruments, Tujunga, CA, USA). For the CCI group, a midline scalp incision was made and a 0.4 cm (ML) × 0.5 cm (AP) craniectomy was performed over the right parietal cortex, leaving the dura intact. CCI was induced using a stereotaxic impactor (Impact One™, Leica Biosystems Richmond Inc., Richmond, IL, USA) to induce mild-to-moderate TBI (3.0-mm tip-oriented perpendicular to the cortical surface, 4.4 m/s velocity, 1.0 mm depth, and 100 ms duration). The bone flap was replaced with dental acrylic (leaving a small hole at the lateral edge for intracranial fluid-pressure relief) and the incision was sutured. Body temperature was maintained at 37 °C throughout the surgery. Respiration rate was visually monitored. Sham animals underwent the same procedures except the CCI impact. Lidocaine (2% solution, topical at incision), Burprenorphine ER (0.6 mg/kg, s.q.), carprofen (5 mg/kg, s.q.), and saline (0.9% solution, 0.5 mL, s.q.) were given for analgesia and fluid replacement. Rats recovered in a temperature-controlled chamber (30 °C ambient air) and monitored for neurological complications. None of the rats exhibited any symptom.

### Hyperpolarization of [1-^13^C]Pyruvate and [2-^13^C]Pyruvate

A 26 μL of neat [1-^13^C]pyruvic acid or [2-^13^C]pyruvic acid sample containing 30-mM AH111501 was polarized using a dynamic nuclear polarization polarizer (SpinAligner, Polarize ApS, Frederiksberg, Denmark) operating at 6.7 T and ∼1.3 K. Each sample was polarized by irradiating microwave (187.995 GHz) for ∼1.5 h until the solid-state polarization build-up curve reached a plateau, then rapidly dissolved in 4.5 mL of pre-heated dissolution medium (D_2_O) and immediately mixed with neutralization solution (0.72 M NaOH, 0.4 M Trizma base, and 0.1 g/L Na_2_EDTA) to achieve a pH of 7.0-7.5. This procedure yielded ∼4.5 mL of HP solution at a final pyruvate concentration of 80 mM and physiological pH. The HP solution was administered intravenously to the rats via the lateral tail vein at a dose of 1.0 mmol/kg body weight and at a rate of 0.25 mL/s.

### *In Vivo* MR Experiment

Rats underwent two sessions of a ^1^H/^13^C-integrated MR protocol, separated by 1 day. Each session included bolus injection of HP pyruvate, and separate groups of animals were used for [1-^13^C]pyruvate and [2-^13^C]pyruvate. In the first group, CCI rats (n = 6) and age-matched controls (n = 5) were examined using HP [1-^13^C]pyruvate at 3 days post-surgery under fed conditions and again at 4 days post-surgery after 20-24 h of fasting. The same two-day MR sessions were performed to the second group (CCI, n = 6; control, n = 5) using HP [2-^13^C]pyruvate. The selected time points were based on prior evidence showing that hepatic APR signaling, particularly MCP-1 (CCL2), peaked at 3 days after TBI.^28^

For each imaging session, rats were anesthetized with isoflurane (2-3%) and cannulated with a tail vein catheter, then positioned tail-first in a supine position in a 3T preclinical MR scanner (Bruker Biospec, Billerica, MA, USA). A dual-tuned ^1^H/^13^C surface coil (20-mm diameter; Bruker) was positioned directly over the liver for both RF excitation and signal reception. Coil placement and animal positioning were verified using a ^1^H gradient-echo localizer (echo time [TE] = 7 ms, repetition time [TR] = 100 ms, field of view [FOV] = 60 × 60 mm^2^, slice thickness = 2 mm). Data acquisition was performed using ParaVision software (version 360.3.4, Bruker). Prior to ^13^C MRS acquisition, localized ^1^H shimming was performed over the liver. HP ^13^C data acquisition was initiated immediately after the start of the pyruvate injection. Time-resolved dynamic ^13^C spectra were acquired using non-localized RF excitation (flip angle = 10°, spectral width/points = 10,000 Hz/4,096, TR = 3 s, #repetition = 60). The ^13^C acquisition frequency was calculated from the water ^1^H frequency^29^ and set to 171 ppm and 100 ppm for hyperpolarized [1-^13^C]pyruvate and [2-^13^C]pyruvate, respectively.

### 13C Data Reconstruction and Analysis

^13^C free induction decay signals were processed in MATLAB R2024b (MathWorks, Natick, MA, USA) using the pvmatlab toolbox for handling Bruker raw data. Spectral preprocessing comprised Gaussian apodization (line broadening corresponding to 1.5× the nominal spectral resolution), followed by zero-filling by a factor of two prior to fast Fourier transform. Zero- and first-order phase corrections were then applied to obtain phase-aligned spectra suitable for peak quantification and display.

HP ^13^C metabolites were quantified by integrating the corresponding resonances from time-averaged ^13^C spectra over the first 90 s. Each metabolic product was normalized to the total HP ^13^C products (TP), excluding circulating HP pyruvate and its non-metabolic product, pyruvate-hydrate. TP was calculated by summing [1-^13^C]lactate, [1-^13^C]alanine, and [^13^C]bicarbonate signals for HP [1-^13^C]pyruvate injection, and by summing [2-^13^C]lactate, [2-^13^C]alanine, [2-^13^C]PEP, [5-^13^C]glutamate, and [1-^13^C]acetyl-L-carnitine (ALCAR) for HP [2-^13^C]pyruvate injection. PC-specific small peaks corresponding to malate and aspartate produced from HP [1-^13^C]pyruvate were observed only in a subset of the rats and not included in the quantitative analysis.

### NMR Isotopomer Analysis

To confirm *in vivo* observation, fourteen rats were prepared separately for *ex vivo* tissue analysis: CCI (n = 4), sham (n = 6) and control (n = 4) rats. At 4 days post-surgery with 20-24 h of fasting, rats were anesthetized with isoflurane (2-3%) and cannulated with a tail vein catheter. A solution that contains thermal (non-HP) 80-mM [U-^13^C_3_]pyruvate was injected intravenously as bolus (0.25 mL/s, 1 mmol/kg body weight), and the liver was harvested 2 min after the injection. After opening the abdominal cavity through a midline incision, the liver was carefully exposed and separated from surrounding connective tissue and ligaments. The portal vein and associated vessels were identified, and the liver lobes were gently lifted using forceps while being detached from vascular attachments. The tissue was removed quickly and immediately immersed in liquid nitrogen to be fixed.

Ground liver tissue (3 g) was treated using perchloric acid (15 mL). The mixture was vortexed for 1 min and centrifuged, and the supernatant was transferred to a new tube. The extraction was repeated, and supernatant was neutralized with KOH, centrifuged, and lyophilized. The dried residue was dissolved in 300-μL D_2_O, containing 4.9-mM 4,4-dimethyl-4-silapentane-1-sulfonic acid as a reference, and centrifuged, and supernatant was transferred to an NMR tube.

^13^C NMR spectra were obtained using a 600 MHz vertical bore high resolution NMR spectrometer (Bruker) equipped with a 3-mm broadband probe with the observe coil tuned to ^13^C (150 MHz). ^13^C NMR spectra were collected using a 60° RF excitation at 25 °C (spectral width/point = 36,058 Hz/144,226, TR = 2 s). Proton decoupling was performed using a standard zg0pg pulse program (Bruker).

^13^C NMR data were processed and analyzed using TopSpin (Bruker, version 4.5.0). Standard phase and baseline corrections were applied prior to peak integration. Metabolite signals were identified based on their characteristic chemical shifts and ^13^C-^13^C coupling patterns. Glucose labeling was assessed from the [5,6-^13^C_2_]glucose doublet observed at the C6 resonance and from the [4,5-^13^C_2_]glucose doublet at the C5 resonance. Integrated peak areas were normalized to the DSS internal reference and used to compare relative flux through PC/PEPCK versus PDH pathways.^25^ PEP labeling was quantified from the C3 multiplet arising from ^13^C-^13^C couplings with C2 (^1^J_23_) and C1 (^2^J_13_) as well as from the C2 multiplets. Both signals were integrated and normalized to DSS to obtain relative ^13^C PEP abundance. Glutamate labeling was evaluated by integrating the C2 doublet (C2-C3) and the C4 doublet (C4-C5). Peak areas were normalized to DSS and reported as relative ^13^C glutamate signal.

### Statistical Analysis

Statistical analyses were conducted using Prism 10 (GraphPad Software; version 10.2.3). The comparison of *in vivo* HP ^13^C products between the CCI and healthy control groups was performed using an unpaired two-tailed Welch’s t test. Comparisons for *ex vivo* NMR isotopomer analysis among CCI, sham, and control groups were performed using Welch’s one-way ANOVA followed by Dunnett’s T3 post hoc multiple-comparisons testing. *P* < 0.05 was considered statistically significant. Values are reported as mean ± standard deviation.

## RESULTS

### Hepatic PDH Flux and Pyruvate Oxidation Remain Unchanged Following TBI

Under normal fed conditions, hepatic [^13^C]bicarbonate production from HP [1-^13^C]pyruvate is predominantly driven by PDH flux (**Figure 1A**)^24,30^. Following an injection of HP [1-^13^C]pyruvate, products including [1-^13^C]lactate, [1-^13^C]alanine, and [^13^C]bicarbonate were detected (**Figure 1B**). The fraction of HP bicarbonate in total HP ^13^C products (TP), [^13^C]bicarbonate/TP, was 0.054 ± 0.014 for the CCI group and 0.073 ± 0.030 in the control group (*P* = 0.256, **Figure 1C**), suggesting CCI has minimal impacts on PDH flux in the liver. [1-^13^C]Lactate/TP was at comparable levels between the CCI (0.639 ± 0.044) and control (0.631 ± 0.057) groups (*P* = 0.811, **Figure 1D**). [1-^13^C]Alanine/TP was 0.297± 0.045 for CCI rats and 0.243 ± 0.035 for controls (*P* = 0.056, **Figure 1E**).

**Figure 1.**
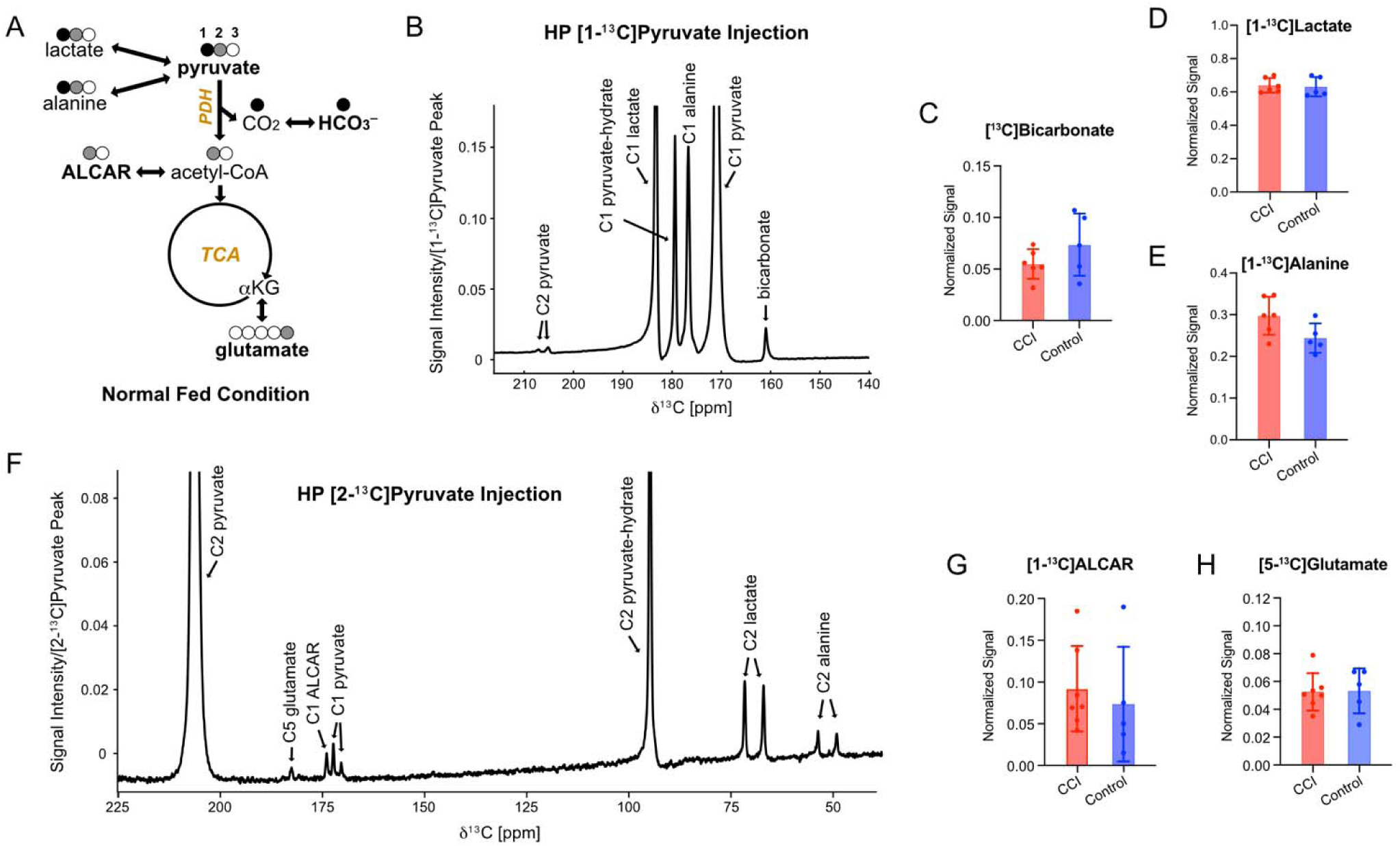
Intact hepatic PDH flux after TBI. (A) HP [1-^13^C]pyruvate detects PDH-mediated decarboxylation via HP [^13^C]bicarbonate production under fed conditions. HP [2-^13^C]pyruvate detects HP [1-^13^C]acetyl-L-carnitine (ALCAR) which makes rapid interconversion with [1-^13^C]acetyl-CoA. [5-^13^C]Glutamate production indicates PDH flux followed by the TCA cycle. The solid circles and open circles represent labeled (^13^C) and unlabeled (^12^C) carbons, respectively. Black and gray circles refer to carbons originated from [1-^13^C]pyruvate and [2-^13^C]pyruvate, respectively. (B) Representative time-accumulated (0-90s) ^13^C spectrum acquired using HP [1-^13^C]pyruvate. (C) HP [^13^C]bicarbonate/TP, (D) [1-^13^C]lactate/TP, and (E) [1-^13^C]alanine/TP were comparable between CCI and control groups. (F) Representative accumulated ^13^C spectrum acquired using HP [2-^13^C]pyruvate. (G) HP [1-^13^C]ALCAR/TP and (H) [5-^13^C]glutamate/TP were comparable between groups. Data are presented as mean ± SD with individual animal values overlaid. Total product (TP) is the sum of [^13^C]bicarbonate, [1-^13^C]lactate, and [1-^13^C]alanine for HP [1-^13^C]pyruvate and the sum of [5-^13^C]glutamate, [1-^13^C]ALCAR, [2-^13^C]PEP, [2-^13^C]lactate, and [2-^13^C]alanine for HP [2-^13^C]pyruvate.

To assess pyruvate utilization through mitochondrial pathways, *in vivo* HP [2-^13^C]pyruvate MRS was performed under the same experimental conditions (**Figure 1F**). Since the labeled carbon is retained in [1-^13^C]acetyl-CoA after PDH (**Figure 1A**), [1-^13^C]ALCAR that makes rapid equilibrium with [1-^13^C]acetyl-CoA was detected at comparable levels between groups under fed conditions, confirming the [^13^C]bicarbonate observation from HP [1-^13^C]pyruvate (**Figure 1G**). Moreover, the level of [5-^13^C]glutamate signal remained unchanged after CCI, indicating hepatic pyruvate oxidation through the TCA cycle was unaltered after TBI (**Figure 1H**). Natural-abundance C2 and C1 signals appeared as ^13^C-^13^C-coupled doublets following HP [1-^13^C]pyruvate and HP [2-^13^C]pyruvate administration, respectively.

### HP Pyruvate Detects Enhanced Hepatic PC-PEPCK Pathway *In Vivo*

Under fasted condition where PDH flux is suppressed, [^13^C]bicarbonate production from HP [1-^13^C]pyruvate reflects the PEPCK-mediated decarboxylation of ^13^C following pyruvate carboxylation (**Figure 2A**).^24–26^ After 20-24 h of fasting, overall [^13^C]bicarbonate production was lower than that measured under fed conditions, confirming fasting-induced PDH suppression (**Figure 2B**). PC-specific downstream products such as aspartate and malate appeared but were not quantified due to inconsistent detection and limited chemical shift dispersion at 3T. [^13^C]Bicarbonate/TP was higher in the CCI group than in the control group (bicarbonate/TP = 0.015 ± 0.007 *vs*. 0.007 ± 0.001, *P* = 0.038; **Figure 2C**). The elevated bicarbonate production in the CCI group under fasting conditions suggests that pyruvate flux through PC-PEPCK is enhanced while pyruvate oxidation is preserved during acute post-TBI. [1-^13^C]Lactate and [1-^13^C]alanine levels showed no differences between the groups (**Figure 2D, E**).

**Figure 2.**
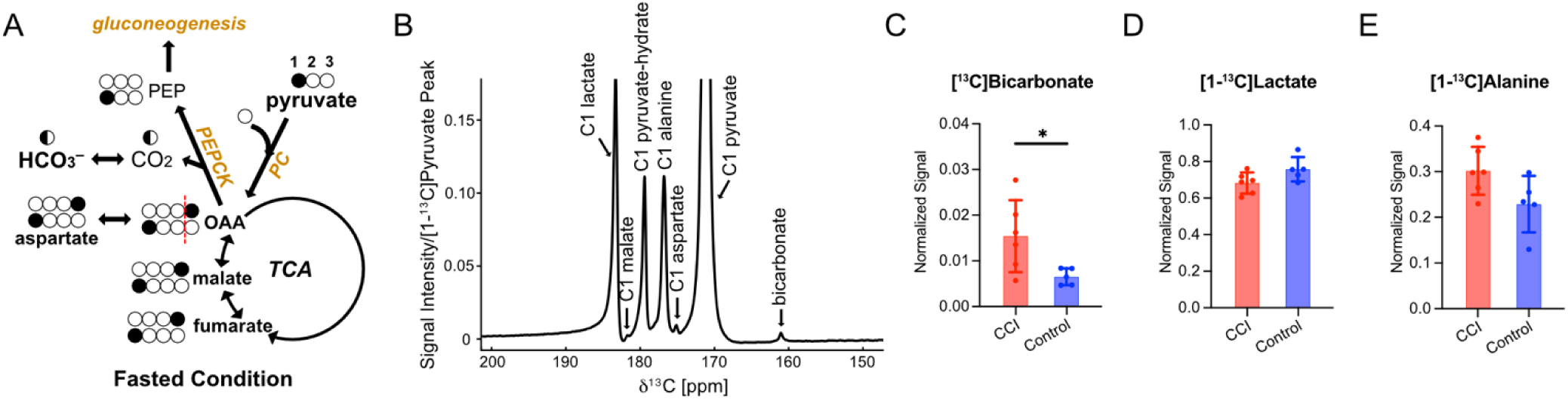
Increased hepatic [^13^C]bicarbonate production in CCI under fasting conditions. (A) Schematic illustration of HP [1-^13^C]pyruvate metabolism through PC-PEPCK pathways. Solid and empty circles indicate ^13^C-labeled and unlabeled (^12^C) carbons, respectively. (B) Representative time-accumulated (0-90s) ^13^C spectrum acquired from the liver following HP [1-^13^C]pyruvate injection under fasted conditions. (C) [^13^C]Bicarbonate/TP increased four days after CCI (*P* = 0.038). Changes associated with CCI were not significant in (D) [1-^13^C]lactate/TP and (E) [1-^13^C]alanine/TP. Data are presented as mean ± SD with each dot representing one animal. Total product (TP) is the sum of [^13^C]bicarbonate, [1-^13^C]lactate, and [1-^13^C]alanine. * indicates *P* < 0.05 with an unpaired two-tailed Welch’s t test.

The findings from HP [2-^13^C]pyruvate were consistent with HP [1-^13^C]pyruvate results (**Figure 3**). Under fasted conditions, the HP [2-^13^C]PEP/TP ratio was significantly higher in the CCI group (0.024 ± 0.006) than in controls (0.012 ± 0.006, *P* = 0.025, **Figure 3C**). In contrast, the HP [5-^13^C]glutamate/TP ratio did not differ significantly (CCI: 0.047 ± 0.032, control: 0.041 ± 0.015, *P* = 0.649, **Figure 3D**). [1-^13^C]ALCAR peak was depleted in both groups, reflecting suppressed PDH flux after fasting conditions (**Figure 3E**). As a combined metric that reflects the flux balance between PDH and PC-PEPCK, a ratio of PEP to the sum of glutamate and ALCAR was calculated using HP [2-^13^C]PEP as a readout of PC-PEPCK flux, and HP [5-^13^C]glutamate and [1-^13^C]ALCAR as a readout of PDH activity. The resulting HP PEP/(glutamate+ ALCAR) ratio was higher in the CCI group than controls (0.602 ± 0.269 vs. 0.278 ± 0.170, *P* = 0.048), supporting the enhanced pyruvate flux through PC-PEPCK (**Figure 3F**).

**Figure 3.**
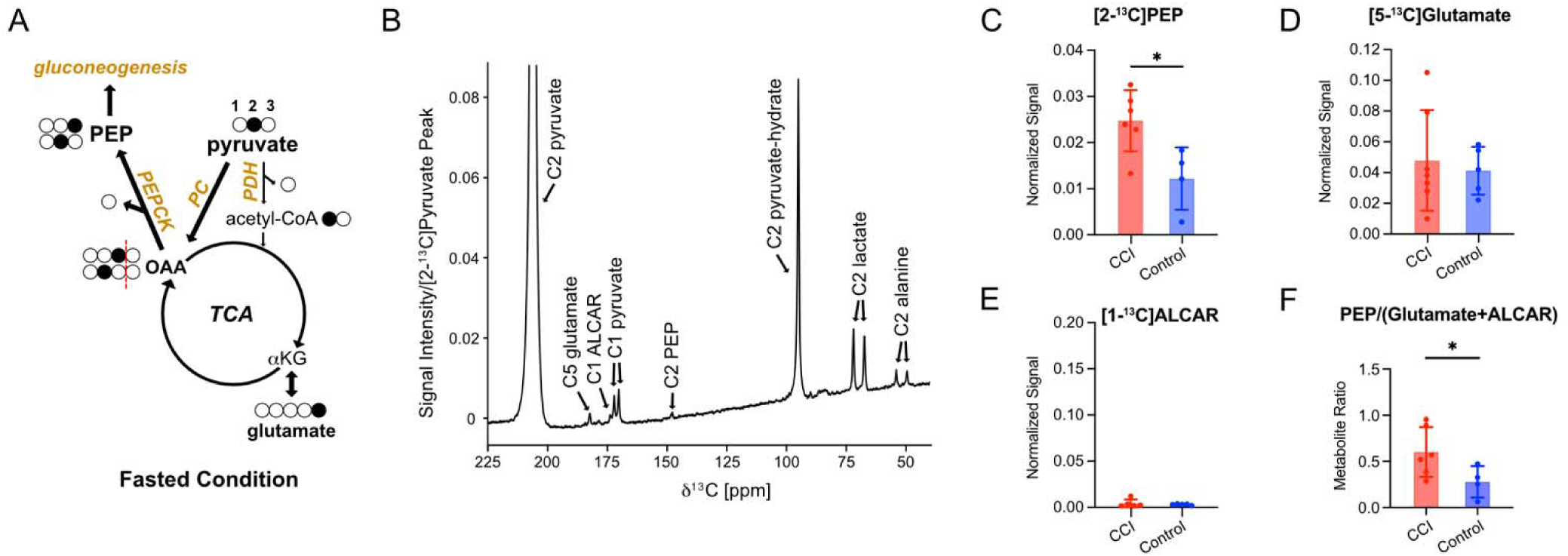
Enhanced [2-^13^C]PEP production in CCI under fasting conditions. (A) Schematic illustration of HP [2-^13^C]pyruvate metabolism under fasted state, highlighting enhanced PC-PEPCK and suppressed PDH pathways. (B) Representative time-accumulated (0-90 s) ^13^C spectrum acquired from liver, under fasted state, following HP [2-^13^C]pyruvate injection. (C) [2-^13^C]PEP/TP increased four days after CCI (*P* = 0.025). (D) [5-^13^C]Glutamate/TP was comparable CCI and control groups. (E) [1-^13^C]acetyl-L-carnitine (ALCAR) was nearly depleted in fasted state. (F) The ratio of PC-PEPCK flux to PDH flux, measured by [2-^13^C]PEP-to-([5-^13^C]glutamate+[1-^13^C]ALCAR), was elevated after CCI (*P* = 0.048). Data are presented as mean ± SD with each dot representing one animal. Total product (TP) is the sum of [5-^13^C]glutamate, [1-^13^C]ALCAR, [2-^13^C]PEP, [2-^13^C]lactate, and [2-^13^C]alanine. * indicates *P* < 0.05 with an unpaired two-tailed Welch’s t test.

### TBI Alters the Hepatic PC/PDH Flux Balance

Liver tissues from the TBI, sham, and control groups were investigated *ex vivo* with NMR following a bolus injection of [U-^13^C_3_]pyruvate to validate the *in vivo* findings. Isotope-labeling patterns of glutamate, PEP, and glucose were analyzed considering three metabolic pathways in mitochondria. First, [U-^13^C_3_]pyruvate can be converted to [1,2,3-^13^C_3_]oxaloacetate and [2,3,4-^13^C_3_]oxaloacetate via PC followed by isotopic (backward) scrambling among four-carbon intermediates (e.g., fumarate) in the TCA cycle. These oxaloacetate isotopomers are decarboxylated by PEPCK to form [1,2,3-^13^C_3_]PEP and [2,3-^13^C_2_]PEP, which ultimately yield [4,5,6-^13^C_3_]glucose and [5,6-^13^C_2_]glucose via gluconeogenesis (**Figure 4A**). Alternatively, the PC-derived oxaloacetate isotopomers can be metabolized through the TCA cycle, producing [1,2,3-^13^C_3_] and [2,3-^13^C_2_]α-ketoglutarate. The α-ketoglutarate isotopomers are either converted into [1,2,3-^13^C_3_] and [2,3-^13^C_2_]glutamate or further metabolized via the TCA cycle to form [1,2-^13^C_2_] and [3,4-^13^C_2_]oxaloacetate. Downstream processing by PEPCK and gluconeogenesis generates [1,2-^13^C_2_]PEP and [3-^13^C_1_]PEP, and subsequently [4,5-^13^C_2_] and [6-^13^C_1_]glucose (**Figure 4B**). The third pathway is via PDH through which [U-^13^C_3_]pyruvate can generate [4,5-^13^C_2_]glutamate, [1,2-^13^C_2_] and [3,4-^13^C_2_]oxaloacetate, and [4,5-^13^C_2_] and [6-^13^C_1_]glucose (**Figure 4C**).

**Figure 4.**
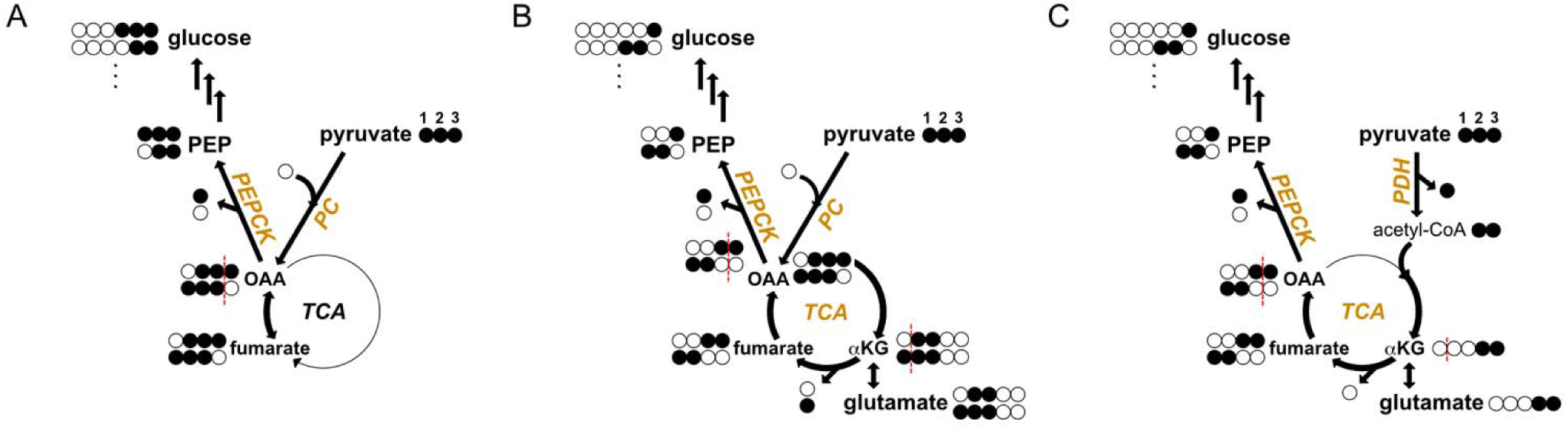
^13^C labeling patterns in glucose, PEP, and glutamate originated from [U-^13^C_3_]pyruvate. (A) PC-mediated pyruvate conversion to oxaloacetate (OAA) followed by a backward scrambling and PEPCK flux can generate [U-^13^C_3_]PEP and [2,3-^13^C_2_]PEP, which can be further metabolized to [4,5,6-^13^C_3_]glucose and [5,6-^13^C_2_]glucose. (B) PC flux followed by forward TCA cycle activity generates different labeling patterns in OAA, labeling [3-^13^C_1_]PEP and [1,2-^13^C_2_]PEP, which can synthesize [6-^13^C_1_]glucose and [4,5-^13^C_2_]glucose via gluconeogenesis, respectively. This pathway labels α-ketoglutarate (αKG) in the TCA cycle, producing [2,3-^13^C_2_]glutamate and [1,2,3-^13^C_3_]glutamate. (C) PDH flux produces distinct labeling in glutamate ([4,5-^13^C_2_]glutamate), but PEP and glucose labeling patterns are the same as the PC-TCA-PEPCK pathway shown in (B).

Based on PEP C3 analysis, the sum of [2,3-^13^C_2_]PEP and [U-^13^C_3_]PEP showed a significant overall difference among the groups (*P* = 0.011) with higher values in the CCI (6.536 ± 1.267, *P* = 0.055) and sham (9.772 ± 3.874, *P* = 0.034) groups compared to the control group (4.059 ± 0.766; **Figure 5A**). Similar trends were observed when analyzing the PEP C2 resonance (**Figure 5B**); The combined abundance of [2,3-^13^C_2_]PEP and [U-^13^C_3_]PEP was higher in the CCI group (3.014 ± 0.326, *P* = 0.005) and the sham group (4.340 ± 1.439, *P* = 0.019) than in controls (1.764 ± 0.121). CCI and sham animals did not differ significantly (*P* = 0.182). Additional splitting of each peak in the spectrum arises from ^13^C-^31^P scalar coupling. Given that these PEP isotopomers originate from combined PC and PEPCK activities, the elevated PEP labeling in CCI animals suggests enhanced hepatic PC-PEPCK flux post-TBI. However, the elevated PEP labeling in sham animals suggests that this response could be partly attributed to the surgical procedure itself or vascular injury.

**Figure 5.**
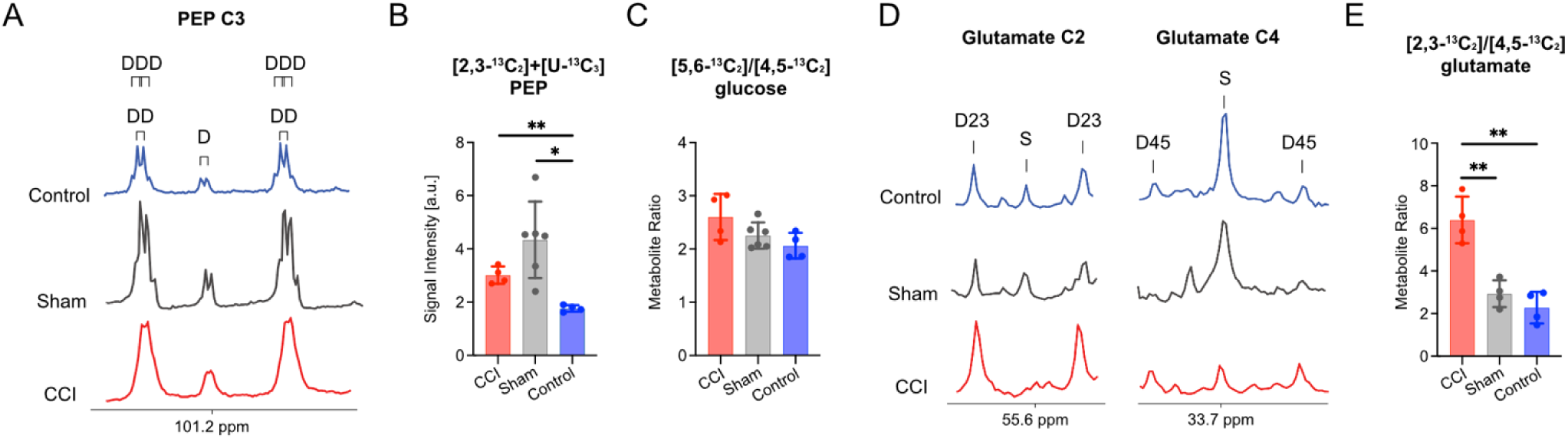
*Ex vivo* ^13^C NMR analysis of liver tissue extracts of CCI, sham, and control animals following bolus injection of [U-^13^C_3_]pyruvate. (A) ^13^C spectral patterns of C3 PEP peaks in control (blue line), sham (black line), and CCI (red line) livers, representing multiplets from [2,3-^13^C_2_]PEP (DD) and [U-^13^C_3_]PEP (DDD). Additional splitting of each peak is due to ^13^C-^31^P coupling. (B) The sum of PEP C2 multiplets were larger in both CCI rats (*P* = 0.005) and sham surgery rats (*P* = 0.019) than in controls, while CCI and sham did not differ (*P* = 0.182). (C) The [5,6-^13^C_2_]glucose-to-[4,5-^13^C_2_]glucose ratio appeared higher in the CCI group, consistent with enhanced PC-PEPCK flux relative to PDH flux, without statistical significance (*P* = 0.186). (D, E) The ratio of doublets in [2,3-^13^C_2_]glutamate (D23) to doublets in [4,5-^13^C_2_]glutamate (D45) was elevated in CCI in comparison to control (*P* = 0.004) and sham animals (*P* = 0.007), indicating the enhanced PC flux relative to PDH flux. Data are presented as mean ± SD. *, ** indicates *P <* 0.05, *P* < 0.01, respectively, using Welch’s one-way ANOVA followed by Dunnett’s T3 post hoc testing.

The ratio of [5,6-^13^C_2_]glucose to [4,5-^13^C_2_]glucose reflects the relative contribution of PC and PEPCK activity compared to PDH and provides an indication of the extent to which pyruvate bypasses the TCA cycle for gluconeogenesis. The [5,6-^13^C_2_]-to-[4,5-^13^C_2_]glucose ratio showed a higher trend in the CCI group (2.603 ± 0.432) than the sham (2.251 ± 0.246) and the control group (2.059 ± 0.241; **Figure 5C**), supporting the observation in PEP C2. However, no significant overall difference was observed among the three groups (*P* = 0.186).

The relative contribution of PC and PDH to TCA cycle entry was evaluated using the ratio of [2,3-^13^C_2_]glutamate to [4,5-^13^C_2_]glutamate, which was significantly higher in the CCI group (6.400 ± 1.099) than both the sham group (2.251 ± 0.246, *P* = 0.007) and controls (2.280 ± 0.742, *P* = 0.004, **Figure 5D, E**), while sham and control animals did not differ (*P* = 0.502). These findings indicate an injury-specific enhancement of PC-mediated anaplerosis into the hepatic TCA cycle following TBI.

## DISCUSSION

In this study, we demonstrated an acute hepatic metabolic response to TBI characterized by increased anaplerotic and gluconeogenic flux. Utilizing *in vivo* MRS with HP [1-^13^C]pyruvate and HP [2-^13^C]pyruvate, we identified an increase in PC-PEPCK pathway after CCI, while markers of PDH-mediated flux remained unaffected. These *in vivo* results were validated by *ex vivo* NMR isotopomer analyses, confirming that central neurotrauma acutely shifts peripheral hepatic metabolism.

The interpretation of HP [^13^C]bicarbonate produced from HP [1-^13^C]pyruvate strongly depends on nutritional state because it can arise from either PDH-mediated decarboxylation or PC/PEPCK-dependent metabolism. Under fed conditions, when hepatic PDH flux predominates, HP [^13^C]bicarbonate levels were similar between TBI and control animals, suggesting preserved pyruvate oxidation through PDH. In contrast, fasting shifts pyruvate utilization toward gluconeogenesis, with PC flux exceeding PDH flux in the liver,^31^ and HP [^13^C]bicarbonate production dominantly reflects PC-PEPCK activity.^24–26^ Therefore, elevated HP bicarbonate in fasted CCI rats indicates enhanced PC-PEPCK flux rather than oxidative pyruvate metabolism.

This interpretation is supported by HP [2-^13^C]pyruvate results, which directly distinguish pyruvate entry into the TCA cycle through PC from oxidative metabolism through PDH. Fasted CCI animals demonstrated significantly increased PEP production *in vivo*, directly indicating enhanced PC-PEPCK activity. In contrast, HP [5-^13^C]glutamate levels, reflecting total entry of pyruvate-derived carbon into the TCA cycle through PDH, were not substantially altered. The elevated HP PEP/(glutamate+ALCAR) ratio observed in the CCI group highlights a shift toward gluconeogenic metabolism relative to oxidative carbon utilization. This conclusion is further supported by isotopomer analysis that showed the increased [2,3-^13^C_2_]glutamate-to-[4,5-^13^C_2_]glutamate ratio in CCI rats compared to sham and control groups, demonstrating preferentially increased PC relative to PDH by brain injury. Collectively, these findings demonstrate that TBI does not increase hepatic pyruvate oxidation through PDH but instead selectively enhances pyruvate flux through the PC-PEPCK pathway to support gluconeogenesis.

The observed hepatic response aligns with the TBI-associated systemic metabolic adaptations. The injured brain experiences extensive alterations in energy metabolism, characterized by accelerated glycolysis, lactate accumulation, and mitochondrial dysfunction.^32^ Our results extend these observations beyond the brain, demonstrating coordinated metabolic communication between the injured brain and the liver.^33^ Upregulation of hepatic gluconeogenesis may be a compensatory mechanism to maintain glucose supply to the injured brain during a period of increased cerebral metabolic demand and impaired brain energy homeostasis.^34^ This interpretation is supported by prior studies reporting TBI-induced systemic hypercatabolic states accompanied by increased hepatic gluconeogenesis.^35–37^ Neuroinflammatory signaling leads to the release of cytokines such as TNF-α, IL-1β, IL-6 and MCP-1, which disrupts hepatic insulin signaling, and promotes glucose production through glycogenolysis and *de novo* gluconeogenesis.^38,39^ Our findings suggest that activation of the PC-PEPCK pathway represents an important metabolic component of this response, contributing to increased endogenous glucose production after TBI.

The sham-operated rats provide valuable insights into the confounding effects of surgical and physiological stress. The large intra-group variability of sham-operated rats and the intermediate level of [5,6-^13^C_2_]glucose-to-[4,5-^13^C_2_]glucose ratio between control and TBI groups suggest that certain hepatic metabolic alterations could be induced by systemic stress responses independent of direct brain injury. In contrast, the [2,3-^13^C_2_]glutamate-to-[4,5-^13^C_2_]glutamate ratio remained at the level of the control group, suggesting that the disrupted balance of PC and PDH is more strongly associated with brain injury rather than procedural stress. We speculate that moderate activation of the hypothalamic-pituitary-adrenal axis following sham surgery may stimulate PC/PEPCK-mediated PEP formation without proportional increase in downstream gluconeogenic flux.^40,41^ For instance, fructose-1,6-bisphosphatase and glucose-6-phosphatase may be control points that limit the conversion of PEP to glucose, resulting in transient PEP accumulation.^42^ In contrast, the robust hormonal and metabolic response induced by TBI may promote gluconeogenesis with increased downstream utilization of PEP toward glucose production, thereby reducing detectable PEP despite elevated PC-PEPCK flux.^43,44^ Future studies incorporating additional measurements of stress hormones and inflammatory mediators are required to precisely distinguish the relative contributions of surgical stress and injury-specific signaling pathways.

The hepatic response identified here may initially be beneficial by maintaining circulating glucose availability for the energy-demanding injured brain. However, sustained activation of gluconeogenic pathways could ultimately become maladaptive, causing hyperglycemia, metabolic stress, and impaired recovery. Clinically, persistent hypermetabolism and dysregulated glucose homeostasis following TBI have been associated with increased morbidity, prolonged hospitalization, and attenuated neurological recovery.^14,45,46^ Therefore, the ability to noninvasively monitor hepatic metabolic fluxes may provide critical insight into the evolution of systemic metabolic dysfunction after brain injury and guide targeted nutritional or metabolic interventions. Our findings suggest that hepatic metabolism as well as systemic glycemic control may be strategic targets to improve metabolic stability and cerebral energy support after TBI.

Several limitations were identified in the current study. First, metabolic measurements were performed at a single post-injury time point for each nutritional condition, based on the prior report indicating maximum neuroinflammatory responses after injury.^28^ Future longitudinal investigations are required to fully characterize the temporal evolution of these metabolic changes. Second, the inherent sensitivity constraints of HP ^13^C MRS limited the detection of downstream metabolites, potentially obscuring more subtle alterations in metabolic flux. Alternative approaches such as methyl-carbon deuteration in [2-^13^C]pyruvate to prolong *T_1_*,^47^ higher magnetic field strength (e.g., 7T) to increase chemical shift dispersion,^48^ and probes that favor PC over PDH flux (e.g., lactate)^49^ could improve HP signal sensitivity and quantification. Finally, the metabolic variability observed in rats with sham surgery highlights the complexity of distinguishing injury-specific effects from systemic stress responses, indicating the necessity for future studies that integrate metabolic, inflammatory, and endocrine measurements to untangle these confounding factors.

In conclusion, TBI is associated with acute hepatic metabolic reprogramming characterized by increased PC-PEPCK pathway activity. By combining *in vivo* HP ^13^C MRS with *ex vivo* isotopomer analysis, we demonstrate that the liver actively participates in the systemic metabolic response to brain injury. Moreover, our results establish hepatic gluconeogenic pathways as potential biomarkers and therapeutic targets for managing TBI and highlight the utility of ^13^C MRS with HP ^13^C-pyruvate for noninvasive assessment of organ-specific metabolic adaptations following acute neurological injury.

## Acknowledgements

The authors thank the UT Southwestern Neuro-Models Facility for the rat CCI surgery (RRID:SCR_022529).

## List of Abbreviations

ALCAR: acetyl-L-carnitine
ALT: alanine aminotransferase
APR: acute phase response
CCI: controlled cortical impact
FOV: field of view
LDH: lactate dehydrogenase
PC: pyruvate carboxylase
PDH: pyruvate dehydrogenase
PEP: phosphoenolpyruvate
PEPCK: phosphoenolpyruvate carboxykinase
TBI: traumatic brain injury
TE: echo time
TP: total HP ^13^C products
TR: repetition time

